# Neogenin-1 marks myeloid-primed fetal hematopoietic stem cells that undergo progressive lineage-restriction with age

**DOI:** 10.64898/2026.09.01.747091

**Authors:** Allison Banuelos, Michelle Baez, Leyla Yılmaz, Elle Koren-Sedova, Monika Zukowska, Andrew T. Burden, Leah Swartzrock-Willner, Uyen Le, Allison Zhang, Benjamin Ohene-Gambill, Nardin Georgeos, Bowen Zheng, Nicole Womack-Gambrel, Ruby Honjol, Rahul Sinha, Irving L. Weissman

## Abstract

Aged hematopoietic stem cells (HSCs) exhibit a shift from balanced to myeloid-biased differentiation, resulting in reduced lymphoid output and impaired adaptive immunity. The question of whether this lineage bias is established in a subset of HSCs during early development or primarily emerges with aging warrants further investigation. Here, we investigate whether myeloid-biased HSCs (my-HSCs) are established at the fetal liver stage by specifically examining Neogenin-1 (NEO1), a previously defined marker of my-HSCs. We identify two distinct populations of *Hoxb5*^+^ HSCs in the fetal liver: NEO1^+^ and NEO1, with NEO1^+^ HSCs exhibiting transcriptional and functional characteristics consistent with my-HSCs. With age, my-HSC-associated transcriptional programs become increasingly reinforced across the *Hoxb5*^+^ pHSC compartment, with NEO1^+^ cells showing early enrichment of this program and both NEO1^+^ and NEO1^−^ cells acquiring broader myeloid-biased features in aging. These findings suggest that lineage programming can begin early in development and is further shaped by age-related changes, potentially contributing to the functional decline observed in the aging hematopoietic system.

## Introduction

Hematopoietic stem cells (HSCs) maintain the lifelong production of all blood lineages through self-renewal and differentiation^1–4^. It has been proposed that the developmental trajectory of HSCs begins early during embryogenesis, progressing from primitive hematopoiesis in the yolk sac to definitive hematopoiesis, perhaps in the yolk sac and/or in the aorta-gonad-mesonephros (AGM) region, and eventually to the fetal liver (FL), which becomes the primary hematopoietic organ until birth^4–11^. During these developmental transitions, HSCs exhibit significant heterogeneity, which is believed to play a key role in shaping their long-term functional properties and lineage potential^12^.

Aging of the hematopoietic system is marked by an increase in myeloid-biased HSCs (my-HSCs), accompanied by a decline in lymphoid output, leading to impaired adaptive immune function^13–15^. Although this myeloid bias is often associated with aging, recent evidence suggests that lineage biases may be established much earlier, potentially during fetal development^16,17^. Determining when and how HSCs begin to adopt lineage biases that persist into adulthood is important for uncovering the mechanisms that drive hematopoietic aging and dysfunction.

Recent studies highlight the importance of specific markers in distinguishing functionally distinct HSC populations^18–25^. Among these, Neogenin-1 (NEO1) has emerged as a key marker that distinguishes my-HSCs from more balanced, lymphoid-producing populations. We have previously found that *Hoxb5* is a transcription factor expressed in long-term repopulating HSCs. We have used *Hoxb5* as a marker alongside NEO1 to further delineate candidate HSC subpopulations: *Hoxb5*^+^ NEO1 HSCs, which exhibit balanced lymphoid and myeloid output, and *Hoxb5*^+^ NEO1^+^ HSCs, which are predisposed to myeloid differentiation^24^. However, the extent to which NEO1 identifies my-HSCs during fetal development has not been investigated.

Identifying how early NEO1^+^ my-HSCs emerge is essential for deciphering the origins of lineage bias and for determining whether this programming is intrinsic to HSCs or driven by extrinsic developmental cues. Our findings provide insight into how NEO1^+^ my-HSCs develop, persist, and potentially contribute to age-related changes in hematopoiesis. Here, we find that NEO1+ HSCs are primed for myeloid differentiation early in life, and as they age, they undergo progressive transcriptional changes that direct their differentiation potential towards myeloid lineages.

## Results

### The fetal liver contains *Hoxb5*^+^ NEO1^+^ pHSCs

We first analyzed available microarray data from Gene Expression Commons, which indicated that *Hoxb5* and *Neo1* are expressed in embryonic day (E) 14.5 fetal liver HSCs (**Supplementary Fig. 1A**). We next examined the expression of additional genes previously associated with myeloid-biased HSCs and found that several components of the adult my-HSC transcriptional signature are also expressed in fetal liver HSCs (**Supplementary Fig. 1B-H**). These observations suggested that molecular features associated with myeloid-biased HSCs may already be present during fetal development and led us to ask whether NEO1 defines a distinct subset of fetal HSCs. We therefore utilized our *Hoxb5*-tri-mCherry mice, in which long-term HSCs are marked by fluorescent mCherry under the regulation of *Hoxb5*, to determine whether NEO1 could be detected on HSCs. Flow cytometry analysis of E14.5 fetal livers from *Hoxb5-*tri-mCherry embryos revealed that a fraction of *Hoxb5*-mCherry^+^ phenotypic HSCs (pHSCs) co-express NEO1 (**Fig. 1A**). NEO1 was also found to be co-expressed at a higher frequency on *Hoxb5*-mCherry^+^ pHSCs compared to *Hoxb5*-mCherry pHSCs (**Fig. 1B**). To confirm these findings, we performed immunofluorescence on fetal liver sections which identified a subset of cells co-expressing c-KIT, mCherry, and NEO1 (**Fig. 1C**). Because NEO1 is detected on both *Hoxb5*-mCherry^+^ and *Hoxb5*-mCherry^-^ pHSCs, we next sought to characterize whether these two subsets of pHSCs exhibit distinct gene expression signatures despite their shared NEO1 protein expression. Single-cell RNA sequencing (scRNA-seq) revealed that *Hoxb5*-mCherry^+^ pHSCs exhibit a gene expression signature more characteristic of undifferentiated pHSCs, while *Hoxb5*-mCherry^-^ pHSCs display a gene expression profile that suggests they are short-term HSCs (ST-HSCs) (**Fig. 1D**). Further analysis of E14.5 FL cells indicated that several HSCs in the fetal liver co-express *Hoxb5* and *Neo1,* with co-expression of *Neo1* being more prevalent in *Hoxb5*-mCherry^+^ pHSCs than *Hoxb5*-mCherry ST-HSCs (**Fig. 1E-F**).

**Figure 1.**
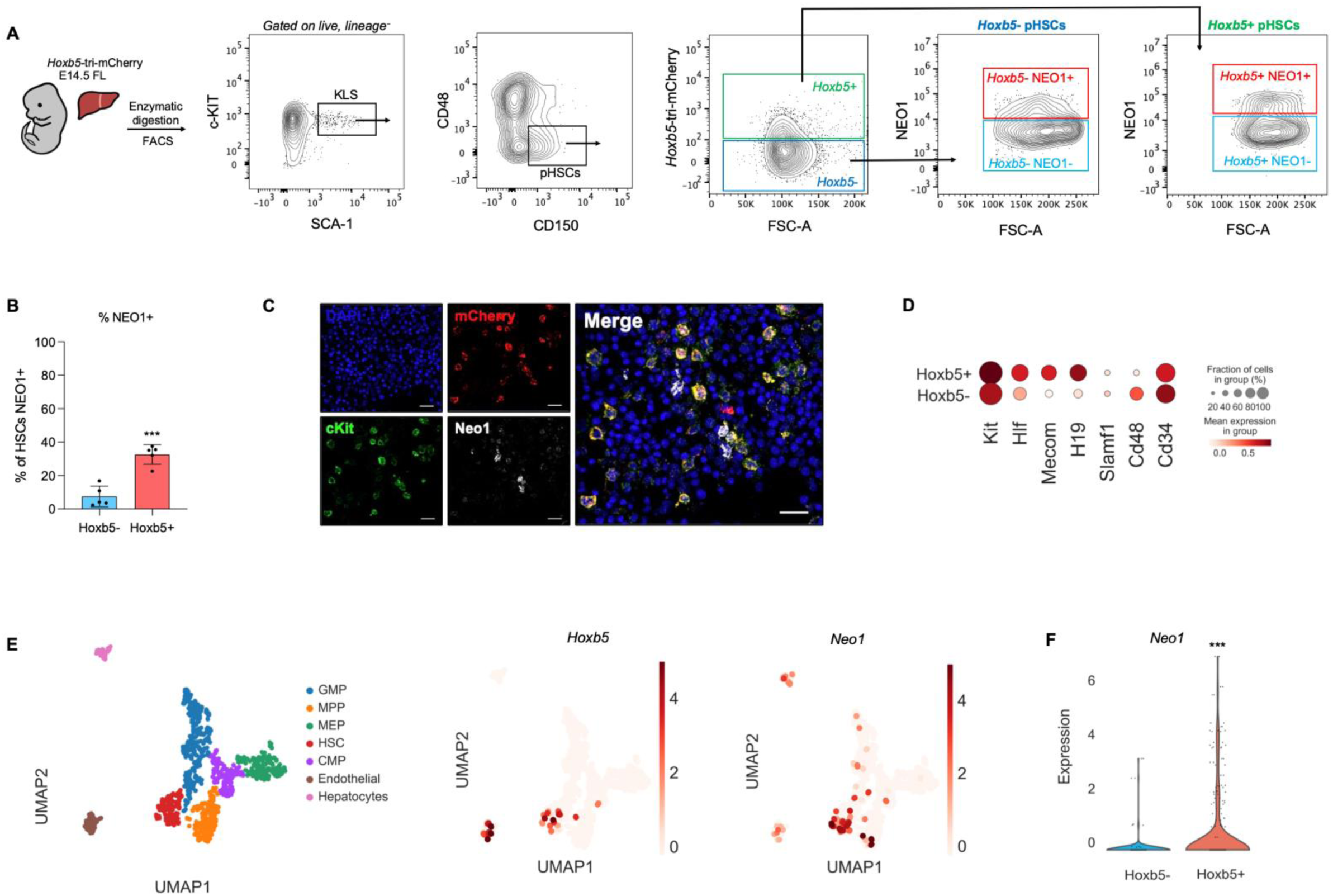
Identification and characterization of *Hoxb5*^+^ NEO1^+^ HSCs in the fetal liver. **(A)** Flow cytometry analysis of *Hoxb5*-mCherry and Neogenin-1 (NEO1) expression in HSCs from E14.5 *Hoxb5*- tri-mCherry mouse fetal livers. Representative flow cytometry plots show the gating strategy used to identify *Hoxb5*-mCherry^+^ and NEO1^+^ HSCs. **(B)** Bar graph showing the frequency of NEO1 on *Hoxb5*- mCherry^+^ and *Hoxb5*-mCherry^-^ HSCs (*n =* 5 embryos). Data are mean ± SEM. \*\*\**P<0.001.* **(C)** Immunofluorescence staining for NEO1, cKIT, and mCherry in an E14.5 fetal liver section from a *Hoxb5*-tri-mCherry reporter mouse (Scale bar = 50 µm). **(D)** scRNA-seq analysis of *Hoxb5*-mCherry^+^ and *Hoxb5*-mCherry^-^ pHSCs from E14.5 fetal liver. Dot plot visualizations show the expression of key hematopoietic stem and progenitor genes in *Hoxb5* -mCherry^+^ and *Hoxb5*-mCherry^-^ pHSCs. The size of the dots represents the percentage of cells expressing the indicated gene, and the color intensity indicates the relative gene expression level. **(E)** scRNA-seq UMAP-embedded Leiden clusters with annotated cell type identities based on expressed transcripts of known genes in the E14.5 FL, and expression plots for *Hoxb5,* and *Neo1*. **(F)** Violin plot of *Neo1 expression* in *Hoxb5*-mCherry^+^and *Hoxb5*-mCherry^-^ pHSCs from E14.5 fetal liver. Pairwise comparisons were performed using the two-sided Mann-Whitney U test. *P<0.05,* \*\**P<0.01,* \*\*\**P<0.001,* \*\*\*\**P<0.0001*.

### *Hoxb5*^+^ NEO1^+^ pHSCs exhibit enhanced expression of my-HSC-associated genes

Having established that the fetal liver contains *Hoxb5*-mCherry^+^ pHSCs that can be further separated into NEO1^+^ or NEO1, we next aimed to determine whether these subsets are transcriptionally distinct in their lineage programming. Previous studies have identified several markers of my-HSCs, including CD150, CD41, CD61, and CD62p^26–39^. We hypothesized that if *Hoxb5*-mCherry^+^ NEO1^+^ pHSCs are myeloid-biased at the fetal liver stage, they may show a coordinated enrichment for transcriptional markers associated with my-HSCs at the single-cell level.

To test this, we performed scRNA-seq on sorted *Hoxb5*^+^ NEO1^+^ and NEO1 pHSCs. For conciseness, we will refer to *Hoxb5*-mCherry^+^ NEO1^+^ and *Hoxb5*-mCherry^+^ NEO1 pHSCs as NEO1^+^ or NEO1 from here on. UMAP demonstrated partial overlap between the two populations, indicating that both subsets occupy related transcriptional space rather than forming completely discrete clusters (**Fig. 2A**). Several genes previously associated with my-HSC identity, including *Esam*, *Eng*, *Procr*, *Tek*, *Selp*, *Itgav*, and *Slamf1*, were preferentially expressed within cells from the NEO1 fraction (**Fig. 2B**). Violin plot analysis confirmed significantly higher expression of multiple my-HSC-associated genes in NEO1 pHSCs relative to NEO1 cells, whereas *Itga2b* did not differ significantly between groups (**Fig. 2C**).

**Figure 2.**
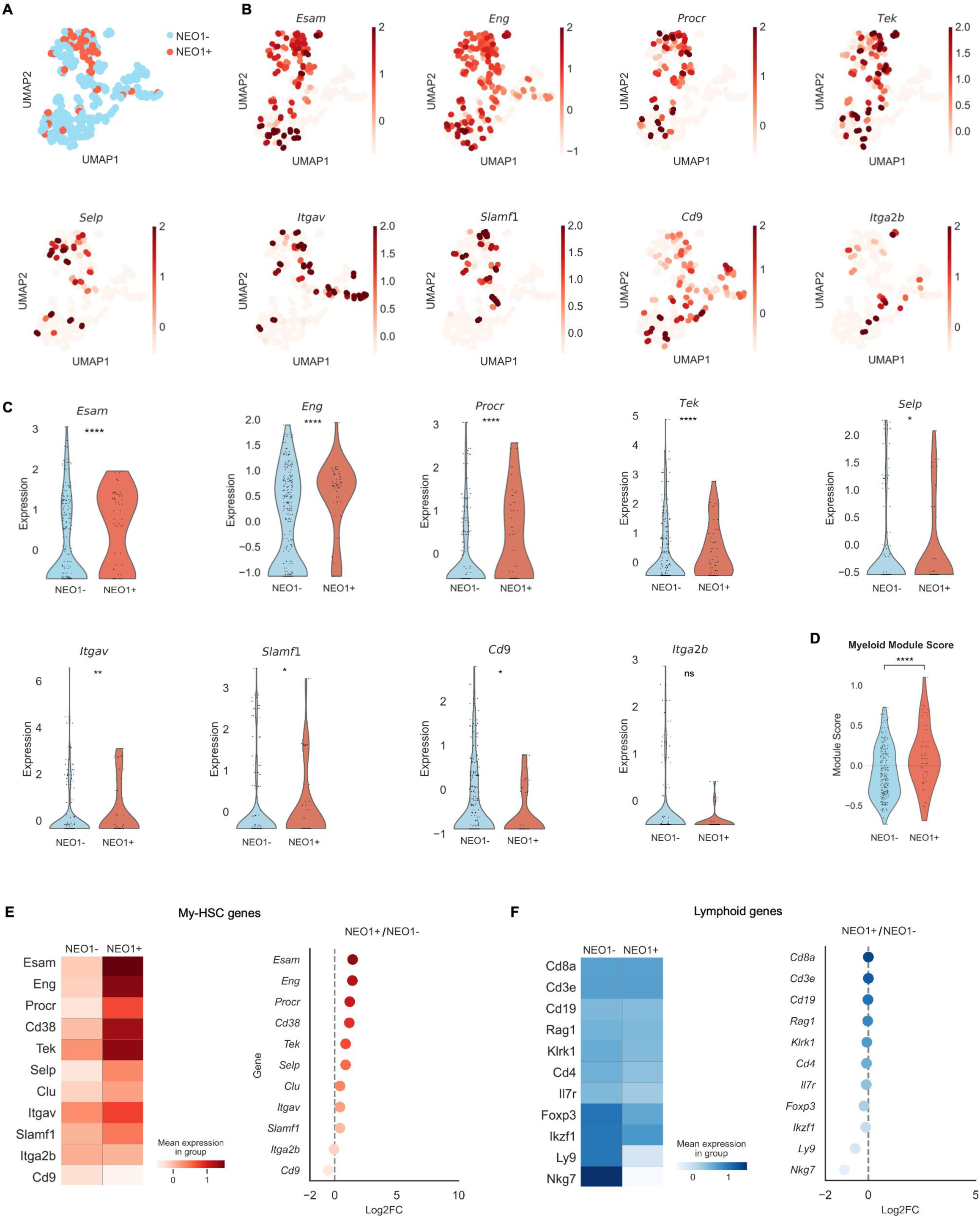
*Hoxb5*-mCherry NEO1 pHSCs are enriched for a my-HSC-associated transcriptional program at the single-cell level. **(A)** UMAP of E14.5 fetal liver Hoxb5-mCherry pHSCs colored by NEO1 status. **(B)** UMAPs of showing expression of selected my-HSC-associated genes (*Esam, Eng, Procr, Tek, Selp, Itgav, Slamf1, Cd9,* and *Itga2b*). **(C)** Violin plots showing expression of selected my-HSC-associated genes in NEO1 and NEO1 E14.5 fetal liver Hoxb5- mCherry pHSCs. **(D)** Myeloid-associated module score calculated from the my-HSC gene set in NEO1 and NEO1 E14.5 pHSCs. **(E)** Summary heatmap of mean expression for my-HSC-associated genes in NEO1 and NEO1 E14.5 fetal liver pHSCs (left) and corresponding mean log2-transformed fold change (log2FC; NEO1 versus NEO1) (right). **(F)** Summary heatmap of mean expression for lymphoid-associated genes in NEO1 and NEO1 E14.5 fetal liver Hoxb5-mCherry pHSCs (left) and corresponding mean log2-transformed fold change (log2FC; NEO1 versus NEO1) (right). Pairwise comparisons were performed using the two-sided Mann-Whitney U test. *P<0.05,* \*\**P<0.01,* \*\*\**P<0.001,* \*\*\*\**P<0.0001*.

To determine whether these gene-level differences reflected a broader coordinated program, we calculated a my-HSC-associated module score using the marker set *Esam*, *Eng*, *Procr*, *Tek*, *Selp*, *Itgav*, *Slamf1*, *Cd9*, and *Itga2b*. NEO1^+^ pHSCs displayed a significantly higher module score than NEO1 pHSCs (**Fig. 2D**), consistent with enrichment of a my-HSC-like transcriptional program within the NEO1 subset. We next examined the average expression of individual my-HSC-associated genes by pseudobulk analysis. This comparison revealed a broader shift toward higher expression of my-HSC-associated genes in NEO1 pHSCs, with the strongest differences observed for *Esam*, *Eng*, *Procr*, *Cd38*, and *Tek* (**Fig. 2E**). Thus, the increased module score in NEO1^+^ cells reflects coordinated differences across multiple components of the my-HSC-associated program rather than being driven by a single gene.

Because my-HSC-associated genes were sufficiently represented across cells to support single-cell and module-level analyses, we focused our primary analyses on this gene set. Additional lineage-associated genes, particularly those associated with lymphoid differentiation, were more sparsely detected **(Supplementary Fig. 2A–B)**. To determine whether enrichment of the my-HSC-associated program in NEO1+ pHSCs was accompanied by a reciprocal lymphoid-associated program in NEO1^−^ pHSCs, we examined a panel of lymphoid-associated genes, including T cell-associated genes (*Cd8a, Cd3e, Cd4,* and *Rag1*), the B cell-associated gene *Cd19*, NK-associated genes (*Nkg7* and *Klrk1*), and broader lymphoid regulators (*Il7r, Foxp3, Ikzf1,* and *Ly9*). Although several of these genes tended to show lower mean expression in NEO1+ pHSCs relative to NEO1^−^ cells, these differences were less pronounced and less uniform than those observed for the my-HSC-associated gene set (**Fig. 2F**). We therefore did not observe evidence of a clear reciprocal lymphoid-biased transcriptional program in the NEO1^−^ fraction.

Together, these data indicate that fetal liver NEO1 pHSCs are more enriched for a my-HSC-associated transcriptional program than NEO1 pHSCs at the single-cell level. Importantly, this distinction appears to be driven primarily by selective upregulation of multiple my-HSC-associated genes rather than broad reciprocal shifts across all lineage-associated markers, and both NEO1 and NEO1 populations retain substantial transcriptional heterogeneity. These findings support the interpretation that NEO1 identifies a fetal liver HSC subset with enhanced my-HSC-like transcriptional features, rather than a fully discrete lineage-committed population.

### *Hoxb5*^+^ NEO1+ pHSCs exhibit a functional bias toward myeloid lineage differentiation in transplantation assays

We next performed transplantation experiments to determine whether there are differences in hematopoietic output between fetal NEO1^+^ and NEO1 pHSCs. We sorted and transplanted 100 NEO1^+^ or NEO1 pHSCs from E14.5 *Hoxb5*-tri-mCherry fetal livers into lethally irradiated adult recipients and monitored hematopoietic output in both primary and secondary transplants (**Supplementary Fig. 3A, Fig. 3A**). Total peripheral blood donor-derived chimerism as determined by CD45.2+ frequency was comparable between recipients of NEO1^+^ and NEO1 pHSCs throughout the course of primary transplant (**Fig. 3B**). NEO1^+^ pHSCs had a significantly higher percentage of myeloid cell output, and a corresponding decrease in B, T, and NK cells among the donor-derived peripheral blood during the first 8 weeks (**Fig. 3C-D**). This early myeloid bias may reflect transient lineage priming that stabilizes over time as the cells engraft and adapt to the adult bone marrow niche. Analysis of bone marrow 16 weeks after primary transplant revealed that recipients of NEO1 pHSCs had significantly higher overall donor chimerism and a greater proportion of donor-derived Hoxb5 NEO1 HSCs compared to the NEO1 group (**Supporting Fig. 3C-D**), suggesting functional and phenotypic stability of the NEO1 subset.

**Figure 3.**
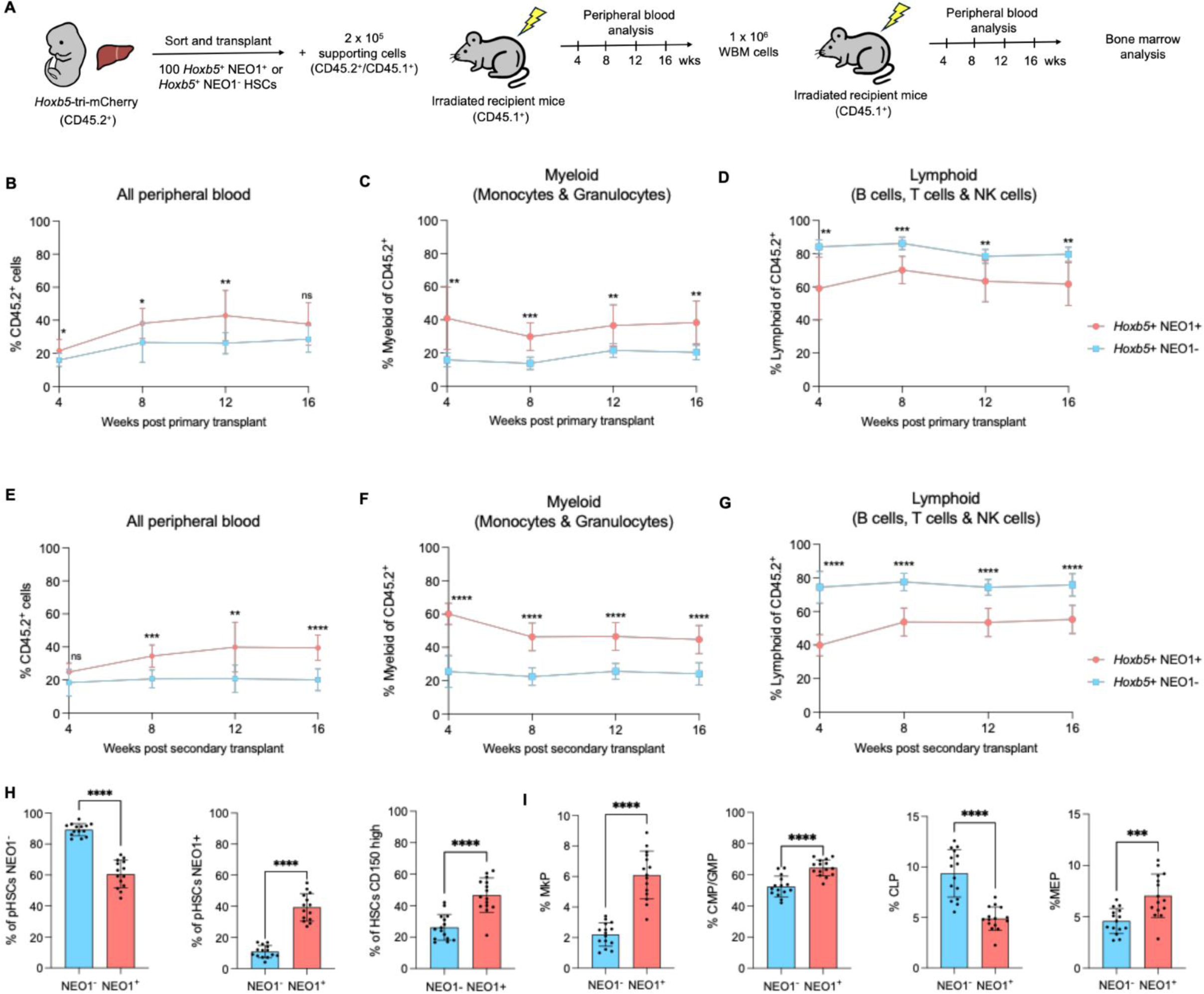
NEO1^+^ fetal liver HSCs exhibit increased myeloid output upon serial transplantation. (A) Schematic diagram depicting the transplantation of E14.5 NEO1^+^ or NEO1 HSCs from the fetal liver of CD45.2^+^ *Hoxb5*-tri-mCherry mice into lethally irradiated CD45.1^+^ recipient mice for primary and secondary transplants. Peripheral blood is evaluated every 4 weeks for 16 weeks following each transplant. At the end of the secondary transplant, recipient bone marrow is analyzed. **(B)** Percent of donor-derived CD45.2^+^ cells among all peripheral blood cells at 4-, 8-, 12-, and 16-weeks post-transplant. **(C)** Percent of myeloid cells (CD11b^+^ GR1^+^ granulocytes and monocytes) among donor-derived CD45.2^+^ cells at 4, 8, 12, and 16 weeks post primary transplant. **(D)** Percent of lymphoid cells (B220^+^ B cells, CD3^+^ T cells, NK1.1^+^ NK cells) among donor-derived CD45.2^+^ cells at 4, 8, 12, and 16 weeks post primary transplant. (“NEO1^+^ recipients”, *n =* 15; “NEO1 recipients”, *n =* 15) **(E-G)** Same as in B-D but analyzing peripheral blood in secondary recipients transplanted with 1×10^6^ whole bone marrow cells from primary hosts **(H-I)** Bone marrow analysis post-secondary transplantation. Bar graphs represent the percentages of donor-derived CD45.2^+^ bone marrow cells including pHSCs that are *Hoxb5*-mCherry^+^ NEO1^+^, *Hoxb5*-mCherry^+^ NEO1 (H), CD150^High^ HSCs, CLPs, GMPs and CMPs combined, MkPs, and MEPs (I) (“NEO1^+^ recipients”, *n =* 15; “NEO1 recipients”, *n =* 15). All graphs in this figure indicate mean ± SEM. \**P<0.05,* \*\**P<0.01,* \*\*\**P<0.001,* \*\*\*\**P<0.0001.* Data shown are derived from 3 independent experiments, with a total of 15 mice across experiments.

To evaluate the long-term reconstitution potential and stability of this lineage output, we performed serial transplantation of whole bone marrow from primary recipients into lethally irradiated secondary recipients. This time, donor chimerism from transplanted NEO1^+^ pHSCs in recipients was significantly higher throughout the transplant process, and they also demonstrated a persistent trend of increased myeloid output throughout the experiment (**Fig. 3E-G**).

Analysis of donor-derived hematopoietic stem and progenitor cells in the bone marrow of secondary recipients at 16-weeks post-transplant revealed that a substantial proportion of the transplanted NEO1^+^ pHSCs retained their NEO1 expression, indicating stability of the marker in a majority of cells (**Supplementary Fig. 3B, Fig. 3H**). However, a subset of these NEO1^+^ cells lost NEO1 expression and converted to NEO1. In contrast, NEO1 HSCs exhibited minimal plasticity, with only a small fraction gaining NEO1 expression post-transplant, while the majority remained NEO1. Further analysis revealed distinct lineage outcomes for NEO1^+^ transplanted pHSCs compared to NEO1. The proportions of CD150^High^ HSCs, common myeloid progenitors (CMPs), granulocyte-monocyte progenitors (GMPs), megakaryocyte progenitors (MkPs), and megakaryocyte-erythroid progenitors (MEPs) were significantly elevated in the NEO1^+^ group relative to NEO1 HSCs (**Fig. 3I**). This was accompanied by a significant decrease in the generation of common lymphoid progenitors (CLPs).

These findings further support the sustained myeloid-biased phenotype of NEO1^+^ pHSCs, underscoring their stable lineage commitment through serial transplantation. These findings indicate that fetal NEO1^+^ pHSCs are functionally biased toward myeloid lineage commitment, whereas NEO1 pHSCs display a more balanced differentiation potential already at the fetal liver stage. The preservation of balanced lineage potential in both primary and secondary transplants of the original NEO1 HSCs suggests that the functional characteristics of myeloid-biased NEO1+ and balanced/lymphoid-biased NEO1 HSCs are maintained. Together with our transcriptional data, these results show that although gene expression differences between NEO1 and NEO1 pHSCs are modest, they are functionally meaningful, with transplantation assays revealing robust and stable lineage biases encoded already at the fetal stage.

### Aging promotes myeloid restriction of *Hoxb5* NEO1^+^ pHSCs

Building on our findings that fetal NEO1^+^ pHSCs share similarities with their young adult and aged counterparts, we next aimed to investigate the dynamics of *Neo1* expression in Hoxb5+ pHSCs across these developmental stages to gain insights into how their transcriptional profiles change with age. scRNA-seq revealed that *Neo1* is expressed in *Hoxb5*^+^ pHSCs across all age groups, with increased expression observed in aged HSCs, as expected (**Fig. 4A-B**). Flow cytometry analysis of *Hoxb5-* mCherry^+^ pHSCs showed that while the frequency of NEO1^+^ cells is expectedly higher in aged pHSCs compared to young adults, fetal pHSCs also exhibit a higher frequency of NEO1^+^ cells than young adult pHSCs (**Fig. 4C**).

**Figure 4.**
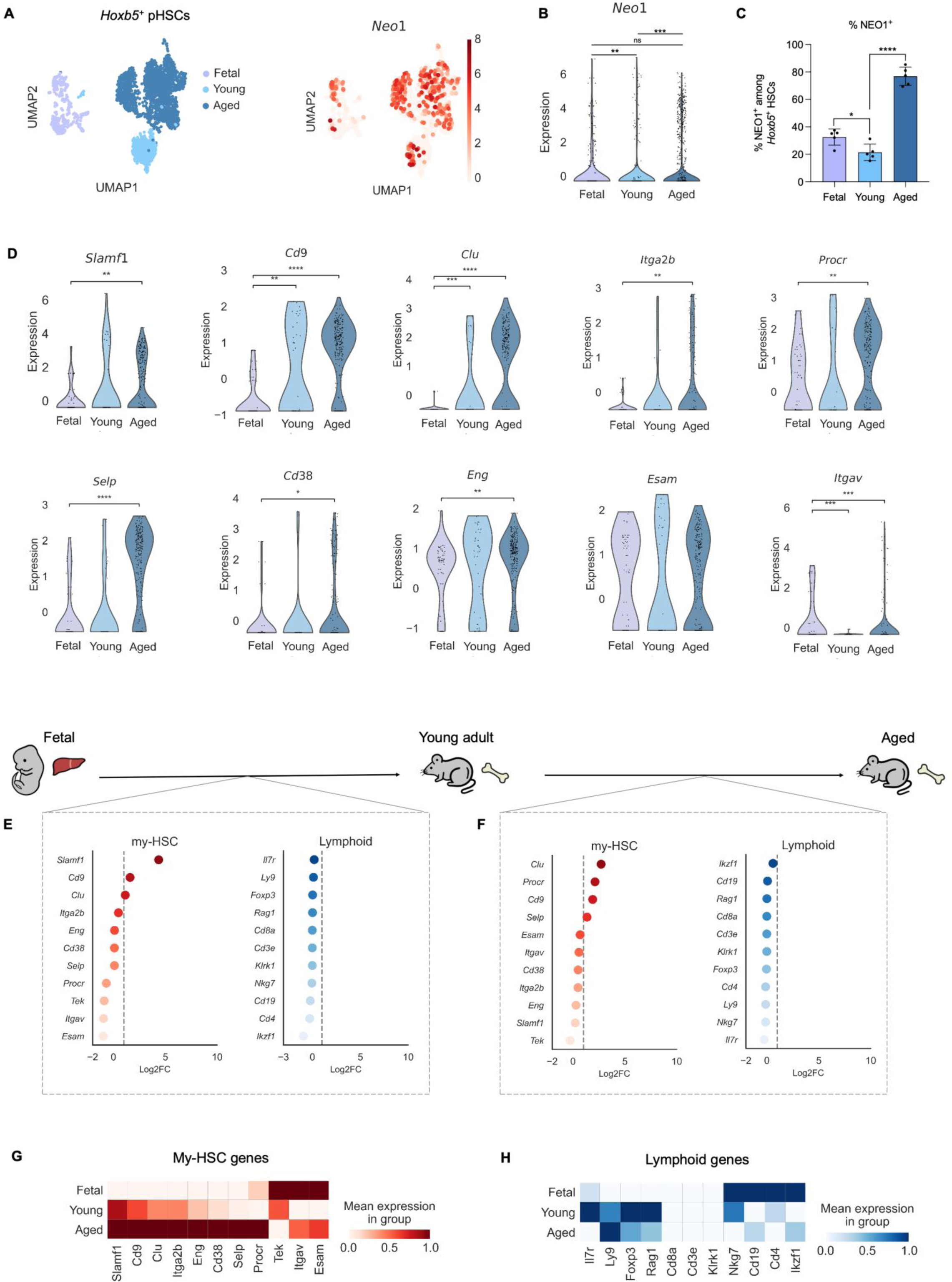
Aging is associated with progressive acquisition of a myeloid-biased transcriptional program in NEO1 pHSCs. **(A)** Plots showing UMAPs of scRNA-seq of fetal (E14.5), young (3 months old), and aged (24 months old) *Hoxb5^+^* HSCs with ages annotated (left) and *Neo1* expression (right). **(B)** Violin plot showing *Neo1* expression in fetal, young, and aged pHSCs. **(C)** Flow cytometry analysis of cell surface NEO1 on *Hoxb5*-mCherry^+^ fetal, young, and aged HSCs (*n* = 5). Error bars indicate mean ± SEM. \**P<0.05,* \*\*\*\**P<0.0001.* **(D)** Violin plots showing expression of representative my-HSC-associated genes (*Slamf1, Cd9, Clu, Itga2b, Procr, Selp, Cd38, Eng, Esam,* and *Itgav*) within NEO1 Hoxb5 pHSCs across fetal, young adult, and aged stages. **(E)** Summary-level comparison of average expression trends for my-HSC- and lymphoid-associated genes in NEO1 pHSCs between fetal and young adult stages. Values are shown as mean log2-transformed fold change (log2FC). **(F)** Summary-level comparison of average expression trends for my-HSC- and lymphoid-associated genes in NEO1 Hoxb5 pHSCs between young adult and aged stages. Values are shown as mean log2- transformed fold change (log2FC). **(G)** Heatmap showing mean expression of my-HSC-associated genes across fetal, young adult, and aged NEO1 Hoxb5 pHSCs. **(H)** Heatmap showing mean expression of lymphoid-associated genes across fetal, young adult, and aged NEO1 Hoxb5 pHSCs. Mean expression values were scaled from 0–1.For panels (C) and (D), pairwise comparisons were performed using the two-sided Mann–Whitney U test. *P<0.05,* \*\**P<0.01,* \*\*\**P<0.001,* \*\*\*\**P<0.0001*.

We next asked whether the NEO1^+^ pHSC compartment exhibits progressive acquisition of a myeloid-biased HSC-like transcriptional program across age. Within NEO1^+^ pHSCs, several canonical my-HSC-associated genes, including *Slamf1, Cd9, Clu, Itga2b, Procr, Selp, Cd38, Eng, Esam,* and *Itgav*, were increasingly enriched with age, consistent with strengthening of a my-HSC-like transcriptional state over time (**Fig. 4D, Supplementary Fig. 4A-L**).

To obtain a broader overview of how this transcriptional state changes across the developmental continuum, we next summarized average expression trends for my-HSC- and lymphoid-associated gene sets within NEO1^+^ pHSCs. At this summary level, the transition from fetal to young adult NEO1+ pHSCs was associated with increased average expression of several my-HSC-associated genes, with *Slamf1* among the most prominent examples, while lymphoid-associated genes showed a modest overall reduction (**Fig. 4E-F**). Comparison of young adult and aged NEO1^+^ pHSCs suggested further reinforcement of the my-HSC-like program, with genes such as *Clu, Procr, Cd9,* and *Selp* showing higher average expression in aged cells (**Fig. 4E-F**). Consistent with this broad trend, summary heatmaps showed a stepwise increase in mean expression of my-HSC-associated genes across fetal, young adult, and aged NEO1+ pHSCs, whereas lymphoid-associated genes showed an overall decline across the same continuum (**Fig. 4G-H**).

Notably, the NEO1− pHSC compartment showed that this age-associated my-HSC-like program is not exclusive to NEO1+ cells (**Supplementary Fig. 5A–D**). While my-HSC-associated genes were minimally expressed in fetal NEO1− pHSCs, they became increasingly detectable in young adult cells and were broadly expressed in aged NEO1− pHSCs. Direct comparison of NEO1^+^ and NEO1^−^ pHSCs across fetal, young adult, and aged stages further illustrated this shift. Differences between the two subsets were most apparent early in life for several my-HSC-associated genes, whereas aged NEO1^+^ and NEO1^−^ pHSCs showed increasingly similar expression patterns **(Supplementary Fig. 5E).** Together, these findings suggest that my-HSC-associated transcriptional features increasingly extend across both NEO1^+^ and NEO1^−^ *Hoxb5*^+^ pHSC compartments with age.

Together, these data suggest that aging broadly promotes acquisition of a myeloid-biased transcriptional program across the *Hoxb5*^+^ pHSC compartment. Although this program is already evident in fetal NEO1^+^ pHSCs, it becomes increasingly represented in both NEO1^+^ and NEO1^−^ pHSCs with age.

### The fetal hematopoietic hierarchy is initiated by *Hoxb5^+^* NEO1^+^ HSCs

In adult hematopoiesis, *Hoxb5* NEO1 HSCs occupy the most primitive position in the hierarchy, as established by previous competitive transplantation studies^24^. To contextualize this known adult organization, we examined pseudotime positioning of adult *Hoxb5* pHSCs and observed the expected arrangement, with Neo1 cells residing in the earliest pseudotime states (**Fig. 5C-D**). We therefore used this adult configuration as a reference point for comparisons across developmental stages.

**Figure 5.**
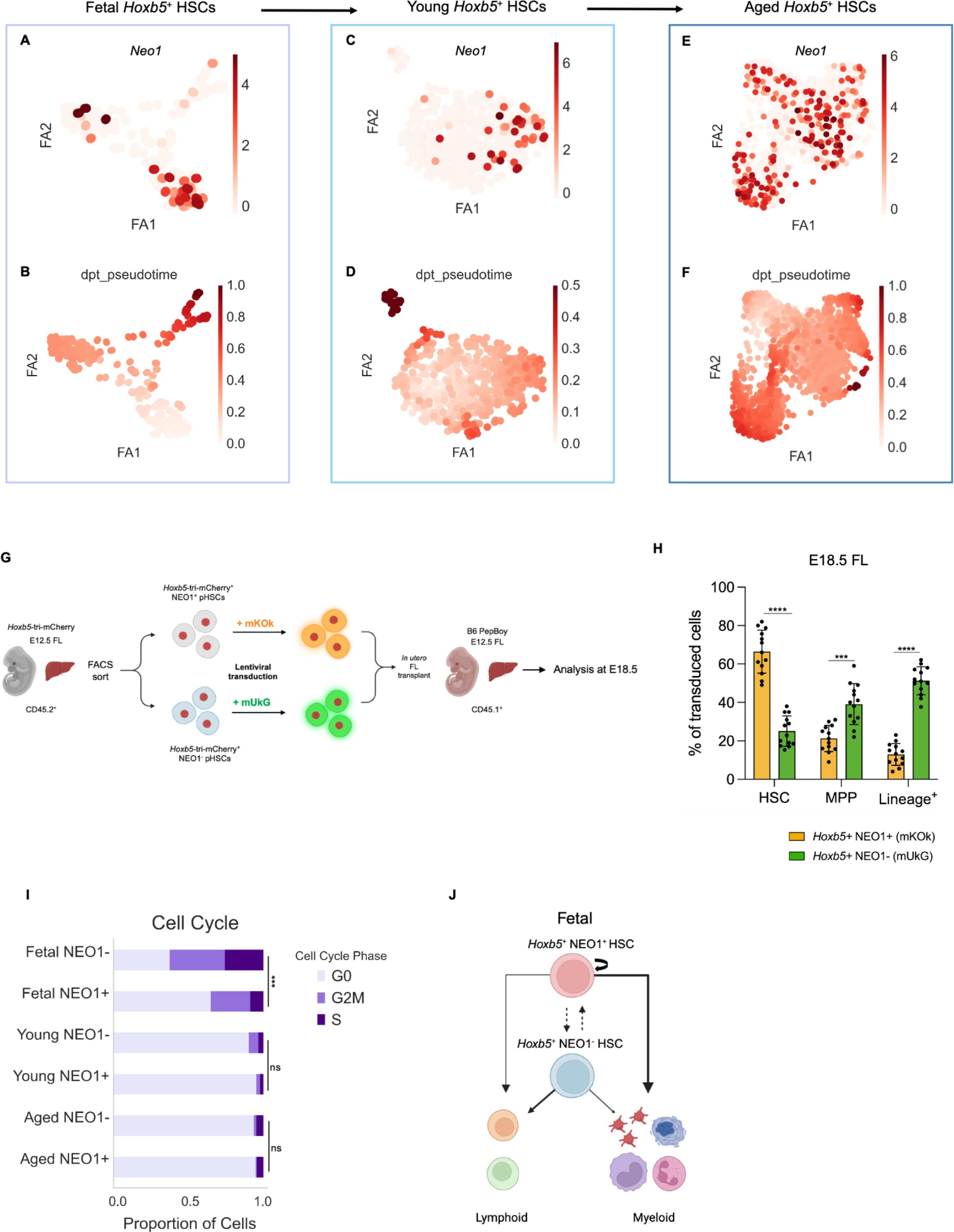
*Hoxb5* NEO1+ pHSCs occupy the earliest identifiable fetal HSC state. **(A-B)** Plot showing PAGA analysis of scRNA-seq data from fetal *Hoxb5^+^*HSCs, with *Neo1* expression overlaid (A) and trajectory analysis showing diffusion pseudotime (dpt) (B). **(C-D)** Plot showing PAGA analysis of scRNA-seq data from young adult *Hoxb5^+^* HSCs, with *Neo1* expression overlaid (C) and trajectory analysis showing diffusion pseudotime (dpt) (D). **(E-F)** Plot showing PAGA analysis of scRNA-seq data from aged *Hoxb5^+^*HSCs, with *Neo1* expression overlaid (E) and trajectory analysis showing diffusion pseudotime (dpt) (F). **(G)** Schematic of *in utero* fetal liver transplantation strategy: *Hoxb5^+^* NEO1^+^ and NEO1 fetal liver HSCs were sorted, transduced with mKOk or mUkG, and co-injected into fetal livers of E12.5 CD45.1^+^ embryos. **(H)** Flow cytometry analysis of transduced *Hoxb5^+^*NEO1^+^ and NEO1 and their contribution to the HSC, MPP, and lineage^+^ compartments in E18.5 fetal livers. Each dot represents an individual embryo (*n = 13*) from three individual experiments. Bars indicate ± SD. Statistical significance was calculated using a two-tailed paired Student’s t-test. *P* < 0.0001 ****, *P* **<** 0.01**. **(I)** Predicted cell cycle phase of *Hoxb5^+^*NEO1^+^ and NEO1 pHSCs across developmental stages, inferred from scRNA-seq data. **(J)** Schematic model of the fetal hematopoietic hierarchy. *Hoxb5* NEO1 pHSCs occupy the earliest fetal HSC state and preferentially maintain the fetal HSC compartment, while *Hoxb5* NEO1 pHSCs represent a downstream fetal HSC state that contributes more extensively to progenitor output. Arrows indicate self-renewal capacity or lineage progression, and line thickness indicates the degree of contribution. Dashed arrows indicate putative developmental relationships inferred from pseudotime analysis and fetal co-transplantation; direct lineage conversion has not been experimentally demonstrated.

We next asked how *Hoxb5* pHSCs are arranged across developmental stages, and specifically how NEO1 expression associates with early HSC ontogeny. To address this, we examined pseudotime ordering of *Hoxb5* pHSCs across fetal, young adult, and aged stages and determined where NEO1- expressing cells localize along these trajectories. In fetal samples, pseudotime analysis placed NEO1 cells in the earliest Hoxb5 pHSC states, indicating that NEO1 expression is associated with the earliest fetal pseudotime positions (**Fig. 5A–B**). This fetal NEO1^+^ cluster also expressed several my-HSC-associated markers, including *Slamf1*, *Selp*, *Clu*, *Cd38*, *Itga2b*, and *Vwf* (**Supplementary Fig. 6A**). In young adult *Hoxb5*^+^ pHSCs, pseudotime organization differed from the fetal pattern, with NEO1 pHSCs occupying earlier states and NEO1*^+^* pHSCs distributed further along the trajectory (**Fig. 5C-D**). Consistent with this organization, my-HSC-associated genes were enriched in cells occupying later pseudotime states, where NEO1 pHSCs were more prominently represented (**Supplementary Fig. 6B**). In aged *Hoxb5* pHSCs, however, NEO1 expression was broadly distributed across the pseudotime manifold, with no clear separation into discrete NEO1 and NEO1 populations (**Fig. 5E–F**). My-HSC-associated markers were likewise broadly expressed (**Supplementary Fig. 6C**), consistent with widespread acquisition of NEO1-associated, myeloid-biased transcriptional features across the aged Hoxb5 pHSC compartment. This pattern is consistent with prior observations that NEO1 HSCs expand with age, and suggests that aging is associated with broad adoption of NEO1-linked transcriptional features across the Hoxb5 pHSC compartment.

To functionally test the developmental ordering suggested by fetal pseudotime, we performed in utero fetal liver co-transplantation experiments. *Hoxb5* NEO1 and NEO1 pHSCs were isolated from E12.5 fetal livers, transduced with distinct fluorescent reporters (mKOk and mUkG, respectively), and co-injected into E12.5 CD45.1 recipient fetal livers (**Fig. 5G**). At E18.5, NEO1 fetal pHSCs contributed predominantly to the HSC compartment, whereas NEO1 cells generated proportionally more MPP and lineage cells (**Fig. 5H**). These results support the interpretation that NEO1 fetal pHSCs represent an earlier fetal HSC state with greater capacity to maintain the HSC compartment, whereas NEO1 fetal pHSCs are relatively more biased toward downstream progenitor output.

Cell cycle analysis further distinguished these fetal pHSC states. Relative to adult Hoxb5 pHSCs, both fetal NEO1 and NEO1 populations exhibited greater proliferative activity overall. However, within the fetal compartment, NEO1 pHSCs were significantly enriched in G0, whereas NEO1 pHSCs showed a relative increase in cycling phases (G2/M and S) (**Fig. 5I**). This relative quiescence of fetal NEO1 pHSCs is consistent with their preferential maintenance of the HSC compartment, while the more actively cycling fetal NEO1 population is consistent with increased progenitor output.

Together, these findings support a model in which *Hoxb5* NEO1 pHSCs represent an earlier functional state during fetal hematopoiesis and preferentially sustain the fetal HSC compartment, whereas NEO1 pHSCs are more proliferative and contribute more extensively to progenitor output (**Fig. 5J**). Although these data establish functional ordering within the fetal compartment, they do not directly demonstrate lineage progression between NEO1 and NEO1 pHSCs. Future lineage-tracing studies will be required to determine whether NEO1 pHSCs can give rise to NEO1 pHSCs in fetal hematopoiesis.

## Discussion

Our previous work has shown that long-term reconstituting *Hoxb5*+ HSCs can be further divided into NEO1^+^ my-HSCs, which favor myeloid differentiation, and NEO1 balanced HSCs (bal-HSCs), which maintain a broader lineage potential^24^. Importantly, NEO1^+^ my-HSCs are already present in adulthood and expand in frequency with age. Here, we extend these findings to examine whether NEO1 similarly identifies my-HSCs at the fetal liver stage and investigate the degree to which myeloid-lineage bias is established early in development.

Our results indicate that fetal NEO1^+^ HSCs display key characteristics of my-HSCs, including a myeloid-biased output upon transplantation, and exhibit a transcriptomic signature associated with my-HSCs. While this signature is further upregulated with age, its early presence in fetal NEO1^+^ pHSCs suggests that myeloid bias is imprinted in a fraction of pHSCs at the fetal stage, preceding the age-related expansion observed in adulthood. Previous hierarchical models of adult hematopoiesis have indicated that long-term bal-HSCs precede my-HSCs in the differentiation process^14,40,41^. Specifically regarding NEO1, our prior studies have shown that NEO1 bal-HSCs reside at the top of the adult hematopoietic hierarchy and may precede NEO1^+^ my-HSCs^24^. However, our findings suggest a different developmental trajectory, where NEO1 pHSCs may arise earlier and contribute to the generation of NEO1 HSCs. Supporting this model, our in utero fetal liver co-transplantation experiments revealed that NEO1 HSCs preferentially contributed to the pHSC compartment, while NEO1 pHSCs generated more downstream MPPs and lineage+ cells, consistent with a role for NEO1 cells as upstream progenitors in fetal hematopoiesis. Lacking lineage tracing, the possible obligatory development of NEO1^+^ HSC from NEO1 HSC is not proven in postnatal bone marrow hematopoiesis.

Whether these are developmentally single lineage HSC or always independent subsets, this observation fits within the framework where the innate immune system plays a crucial role as the primary defense mechanism early in life. As the immune system matures, the emergence of balanced HSCs with broader lineage potential allows for the generation of both myeloid and lymphoid lineages, essential for comprehensive immune surveillance, the generation of immunological tolerance of lymphocytes to self, and adaptive responses.^42–45^ The transition from myeloid-primed NEO1^+^ HSCs being dominant during fetal development to balanced NEO1 HSCs taking precedence in adulthood reflects an adaptive reorganization of the hematopoietic system, ensuring the immune system can meet the increasingly complex demands across different life stages.

As individuals age, a resurgence of myeloid-biased HSCs occurs, contributing to the chronic inflammatory state observed in the elderly, increasing susceptibility to infections and age-related diseases^46–51^. This suggests that the dynamic nature of the hematopoietic hierarchy not only reflects the changing immunological demands throughout life, but also suggests the influence of extrinsic factors, such as alterations in the bone marrow niche, systemic inflammation, and aging-associated senescence. These environmental changes can further exacerbate the myeloid skew by impacting cytokine signaling, niche cell interactions, and extracellular matrix components, creating a feedback loop that reinforces myeloid bias and impairs balanced hematopoiesis^22,38,51,52,53^. This suggests that while intrinsic genetic programs within HSCs establish lineage biases, extrinsic cues from the surrounding microenvironment may play a pivotal role in modulating these biases, particularly during aging. Another consideration is that these aged HSCs may arise from either the cell-intrinsic transition of balanced to myeloid-biased LT-HSCs or the clonal expansion of preexisting fractions of myeloid-biased LT-HSCs, driven by the loss of function of epigenetic modifiers such as TET2 and DMNT3A^54–57^.

Understanding the early establishment of this bias also sheds light on potential intervention points to counteract age-related immune decline. We have recently demonstrated that depletion of my-HSCs, such as those expressing cell-surface markers like NEO1, CD150, or CD62p can rejuvenate the aged immune system^58^. This current study provides insight into the early events that lead to myeloid bias and skewing in aged individuals, starting from the fetal stage. The developmental precedence of NEO1 HSCs raises the possibility that early fetal lineage imprinting may leave lasting epigenetic or transcriptional signatures that influence HSC function throughout life. The identification of NEO1^+^ my-HSCs during fetal development highlights the role these cells play in setting the foundation for myeloid-biased hematopoiesis, which becomes more pronounced with age. By elucidating the early origins of myeloid bias, this study provides critical insights for developing interventions to rejuvenate aged immunity and extend the functional lifespan of the immune system.

### Limitations of the study

While our findings provide novel insights into the early establishment of myeloid bias in fetal NEO1+ HSCs, several key questions remain. First, a full understanding of the hierarchical order and differentiation sequence of NEO1^+^ pHSCs will require additional studies using single cell tracking or lineage tracing experiments. As we have previously demonstrated, yolk sac blood islands not only exist prior to liver hematopoiesis, but synchronic orthotopic transplants of these cells into allogenic recipient yolk sac cavities can give rise to both lymphoid and myeloid blood cells long after birth and during the aging of the recipients^4^. This raises the possibility that the two populations observed in the fetal liver may have originated from yolk sac progenitor HSCs that were either homogeneous or had already bifurcated in their development from mesodermal precursors. Given that no monoclonal antibody separation of yolk sac cells was available in 1975 when those studies were conducted, further definitive and physiological studies will be required to clarify these relationships. These experiments are necessary to trace the fate of individual NEO1^+^ and NEO1 pHSCs over time and to map out their contributions to hematopoiesis across different life stages. Moreover, it remains unclear whether the NEO1^+^ HSCs identified in fetal life persist into adulthood and clonally expand with age, or if adult NEO1^+^ HSCs are derived from a separate population of NEO1 HSCs that acquire NEO1 expression later in life. Distinguishing between these possibilities will be critical to understanding the mechanisms underlying the observed shift towards myeloid bias with aging. Addressing these limitations will not only provide clarity on the developmental trajectory of NEO1^+^ my-HSCs, but will also offer further insights into the potential intervention points for mitigating age-related immune decline.

## Supplementary Figure Legends

**Supplementary Figure 1. Expression of *Hoxb5* and my-HSC markers in fetal, young, and aged HSPCs and HSCs**. **(A)** *Hoxb5,* **(B)** *Neo1 (*NEO1), (**C)** *Slamf1* (CD150), **(D)** *Cd38* (CD38), **(E)** *Itgav* (CD51), **(F)** *Procr* (CD201), **(G)** *Tek* (TIE2), **(H)** *Esam*, **(I)** *Eng* (CD105), **(K)** *Cd9* (CD9). Data and figures from **A-K** were generated and obtained directly from Gene Expression Commons^34^; scale bars represent log2 Signal Intensity and Gene Expression Activity as defined by Gene Expression Commons^34^.

**Supplementary Figure 2. Single-cell analysis of lineage-associated genes in NEO1 and NEO1 pHSCs. (A-B)** Violin plots showing expression of additional lineage-associated genes not displayed as individual single-cell plots in Figure 2, including (A) the my-HSC-associated genes *Cd38* and *Clu* and (B) the lymphoid-associated genes *Cd19, Il7r, Foxp3, Ikzf1, Nkg7,* and *Ly9*. Statistical significance was assessed by two-sided Mann–Whitney U test; ns, not significant; *\*P < 0.05, **P < 0.01, ***P < 0.001, ****P < 0.0001*.

**Supplementary Figure 3. Gating strategy for assessing peripheral blood and bone marrow chimerism of CD45.2+ donor cells following transplantation. (A)** Following exclusion of TER119^+^ red blood cells, donor-derived cells can be distinguished by CD45.2^+^, host cells as CD45.1^+^ and supporting cells as CD45.1^+^CD45.2^+^. Following gating on CD45.2^+^ donor-derived cells, myeloid cells were identified as CD11b^+^ and lymphoid cells were identified as CD3^+^ T cells, B220^+^ B cells, and NK1.1^+^ NK cells. **(B)** Gating strategy for donor-derived HSPCs following secondary transplantation. Following exclusion of Lineage^+^ cells, donor derived bone marrow cells can be distinguished by CD45.2^+^, host cells as CD45.1^+^ and supporting cells as CD45.1^+^CD45.2^+^. **(C)** Percent of CD45.2^+^ donor-derived chimerism in total bone marrow 16 weeks after primary transplant. **(D)** Percent of Hoxb5 NEO1 HSCs in recipient bone marrow after primary transplant, grouped by donor type. Each dot represents an individual mouse (*n = 5*). Bars indicate ± SEM. Statistical significance was calculated using a two-tailed paired Student’s t-test. *P* < 0.001 ***, *P* < 0.01**.

**Supplementary Figure 4. Expression my-HSC markers in fetal, young, and aged *Hoxb5^+^* pHSCs. (A)** UMAP showing scRNA-seq of fetal, young, and aged *Hoxb5*^+^ pHSCs with ages annotated. Gene expression for **(B)** *Slamf1* (CD150), **(C)** *Cd9* (Cd9), **(D)** *Cd38* (CD38), **(E)** *Procr* (CD201), **(F)** *Esam*, **(G)** *Itgav* (CD51), **(H)** *Tek* (TIE2), **(I)** *Eng* (CD105), **(J)** *Clu,* **(K)** *Vwf* and **(L)** *Itga2b*.

**Supplementary Figure 5. Age-associated transcriptional changes in *Hoxb5^+^* NEO1 pHSCs (A)** UMAP showing scRNA-seq of fetal (E14.5), young (3 months old), and aged (24 months old) *Hoxb5^+^ Neo1* pHSCs with ages annotated. **(B)** Gene expression UMAPs of key my-HSC-associated genes (*Slamf1*, *Itga2b*, *Clu*, *Cd38*, and *Selp*) in *Hoxb5^+^ Neo*1 HSCs across developmental stages. **(C-D)** mRNA expression matrix showing the mean expression of my-HSC and lymphoid marker genes in fetal, young, and aged *Hoxb5^+^ Neo1* HSCs. **(E)** Violin plots comparing the expression of selected my-HSC genes (*Slamf1*, *Itga2b*, *Clu*, *Itgav*, *Cd38, Selp, Eng, Esam*) in *Hoxb5*^+^ Neo1 and Neo1 HSCs across fetal, young, and aged stages. Pairwise comparisons were performed using the two-sided Mann-Whitney U test. *P<0.05,* \*\**P<0.01,* \*\*\**P<0.001,* \*\*\*\**P<0.0001*.

**Supplementary Figure 6. Visualization of my-HSC marker dynamics in PAGA-embedded Leiden clusters. (A)** UMAP showing scRNA-seq data for fetal, **(B)** young, and **(C)** aged *Hoxb5*^+^ pHSCs, with PAGA-embedded Leiden clusters. Selected genes *Slamf1*, *Selp*, *Clu*, *Cd38, Itga2b* and *Vwf* are highlighted to demonstrate expression across age groups. Bar plot for each developmental stage showing mean pseudotime value for each Leiden cluster.

## Material and methods

### Mice

*Hoxb5*–tri-mCherry (CD45.2 C57BL/6J background) mice were used as donor cells for transplantation as well as for analysis. Mice were bred at our animal facility according to NIH guidelines. 8-to-12-week old female B6.SJL- Ptprca Pepcb/BoyJ mice (Jackson Laboratory) were used as recipients for transplantation assays. Supporting cells were collected from B6.SJL-Ptprca Pepcb/BoyJ × C57BL/6J (F1 mice CD45.1+/CD45.2+). All animal protocols were approved by the Stanford University Administrative Panel on Laboratory Animal Care.

### Isolation and preparation of fetal liver and bone marrow tissue

Fetal livers were carefully dissected from embryos using fine forceps under a dissection microscope to ensure precision and minimize tissue damage. The dissected liver tissues were then finely minced and incubated at 37C for 30min in pre-warmed digestion media containing M199 media (Thermo), 0.1% Pluronic’s 188 (Thermo), 2mg/mL Colleganse II (Thermo), 0.2mg/mL DNAseI, and 0.1% PVA (Thermo). Liver tissues were then gently passed through a 100 μm filter to remove any debris and clumps of tissue, ensuring a single-cell suspension. Following filtration, the cells were washed twice in PBS with 1% PVA (FACS buffer) to remove any residual contaminants and prepare them for antibody staining.

For young and aged bone marrow isolation, the tibia, femur, and pelvis were carefully dissected from the mice. The bones were then cleaned and crushed using a mortar and pestle in FACS buffer. After crushing, the supernatant containing the cellular components was collected for further processing. Red blood cells were depleted by incubating cells with ACK lysis buffer (Thermo) for 10 mins at room temperature, followed by centrifugation to pellet the remaining cells. Cells were Fc-blocked with TruStain FcX (BioLegend) for 10 minutes prior to staining with antibodies. To isolate c-KIT+ cells, the samples were further incubated with anti-c-KIT APC-eFluor780 (Thermo) for 20 mins at 4C. Following this incubation, the cells were washed twice with FACS buffer. After washing, the cells were incubated with anti-APC beads for 10 mins at 4C to enable the separation of c-KIT+ cells. The c-KIT-enriched cells were then isolated using LS magnetic columns (Miltenyi Biotec) according to the manufacturer’s protocol.

To isolate supporting cells from CD45.1+CD45.2+ “F1” mice, bone marrow was first harvested as previously described. Following isolation, bone marrow cells were incubated with SCA depletion beads (Miltenyi Biotec). The SCA depletion beads were added in accordance with the manufacturer’s instructions. After incubation, the cell suspension was washed and transferred to LS magnetic columns (Miltenyi Biotec) to facilitate the separation of SCA-depleted cells. The flow-through fraction, containing the SCA-1-depleted supporting cells, was collected for transplantation.

### Flow cytometry and cell sorting

Flow cytometry and cell sorting were performed on a FACS Aria II cell sorter (BD Biosciences) or FACS Discover S8 cell sorter (BD Biosciences) and analyzed using FlowJo software. As described previously, we defined HSCs in the fetal liver as: Lineage- Sca1+ Kit+ (KLS) CD150+ CD48-. HSCs in the adult bone marrow defined as: Lineage- Sca1+ Kit+ (KLS) CD150+ FLT3- CD34-. The following antibodies were included in the HSC and progenitor panel: c-Kit (APC-eFluor870), Sca-1 (BV395), CD150 (BV421), CD48 (AF700), CD16/32 (PE), IL7RA (BV510), and CD34 (BV711). Exclusion using the following cocktail of biotinylated “lineage” antibodies were also used to identify HSCs: Ter119, CD4, CD8, B220, Gr1 and NK1.1, followed by secondary staining with BV737-conjugated Streptavidin. NEO1 was detected using a polyclonal primary antibody specific to NEO1 (R&D Systems, Catalog #: AF1079). Samples were washed to remove unbound primary antibody, and secondary staining was performed using a donkey anti-goat Alexa Fluor 488 conjugated antibody. Samples were incubated with anti-mouse CD16/32 Fc Block (BD Biosciences) for 10 minutes prior to staining with antibodies. Antibody staining was performed at 4 °C and cells were incubated for 30 min. Before sorting or analysis, cells were stained with SYTOX Red Dead Cell Stain (Life Technologies) to assess viability as per the manufacturer’s recommendations.

### HSC transplants and peripheral blood analysis

B6.SJL- *Ptprc Pepc^b^*/BoyJ (Jackson Laboratory) recipient mice received a split dose of irradiation (two doses of 4.5Gy, separated by 4 hours) for a total dosage of 9Gy 24h before transplantation. For transplantation assays, 100 sorted donor NEO1+ or NEO1 HSCs (KLS CD150+ CD48- Hoxb5-tri- mCherry+) from *Hoxb5*-tri-mCherry mice were first combined with 2×10^5^ Sca-1 depleted whole bone marrow cells from B6.SJL- Ptprca Pepcb/BoyJ × C57BL/6J F1 mice (CD45.1+/CD45.2+) in 200 μl of PBS with 2% FBS, then injected into the retro-orbital venous plexus. Peripheral blood analyses were performed at 4, 8, 12, and 16 weeks after primary and secondary transplantations. At each time point, 50uL of blood was collected retro-orbitally and added to 500uL PBS with 2mM EDTA. Red blood cells were lysed using ACK lysing buffer (Thermo) for 5 min at room temperature followed by blocking with anti-mouse CD16/32 mouse BD Fc Block (BD Biosciences). Leukocytes were stained with antibodies against CD45.2 (BV421), CD45.1 (BV785), Ter119 (BV737), CD11b (FITC), Gr-1 (BV395), CD8 (PE), CD4 (PE-Cy7), NK1.1 (PE-Cy5), and B220 (APC-Cy7). For each mouse, the percentage of donor chimerism in the peripheral blood was defined as the percentage of CD45.1^-^ CD45.2^+^ cells among total CD45.1−CD45.2+ and CD45.1+CD45.2+ cells.

At the end of secondary transplantation, mice were euthanized and bone marrow from femurs, tibias, and pelvises were prepared as described above. For HSC and progenitor analysis, cells were stained with the following antibodies: CD45.2 (BV421), CD45.1 (BV785), c-KIT (APC-eFluor870), Sca-1 (BV395), CD150 (BV650), FLT3 (PerCP-eFluor710), CD41 (BV510), CD34 (BV711), CD16/32 (PE), IL7Ra (AF700), and NEO1 detected using a polyclonal anti-NEO1 primary antibody followed by donkey anti-goat Alexa Fluor 488 secondary antibody. A lineage cocktail comprising biotin-conjugated antibodies targeting Ter119, CD4, CD8, Gr-1, B220, and NK1.1 was used to exclude differentiated cells. Lineage- positive cells were subsequently stained with BV737-conjugated Streptavidin.

### Tissue staining and confocal imaging

*Hoxb5*-tri-mCherry embryos were dissected at E14.5 under a dissecting microscope using 31 G insulin syringe needles. Fetal livers were embedded in O.C.T. for sectioning and stored at −80C. Before sectioning, fetal livers were fixed in 4% P.F.A. for 35 minutes and washed 3x in PBS with 30 minutes in between each wash. For immunohistochemistry, 7 um fetal livers are placed in 5% donkey serum with 0.5% tritonX and PBS mix for 2 hours at room temperature. Primary antibodies were added to a 1% B.S.A. with 0.5% tritonX and PBS solution at a 1:100 dilution; liver sections were incubated in primary antibodies overnight at 4C. Samples were washed 3x in 0.5% triton X and PBS solution for 30 minutes in between washes before adding secondary antibodies. These antibodies were added to a DAPI with 0.5% tritonX and PBS solution at a 1:250 dilution; liver sections were incubated in secondary antibodies for two hours at room temperature. Samples were washed in 0.5% triton X and PBS solution for 30 minutes before adding RapiClear 1.52 (Sunjin Lab) and incubating the sample overnight. Sections were mounted in Prolong diamond (ThermoFisher) on a superfrost microscope slide and imaged on the Zeiss LM980 Airyscan 2 confocal at 10x-20x magnification. The following primary antibodies were used: anti-mCherry (ab205402), anti-mouse c-KIT (Thermo Fisher 14-1171-82).

### Single-cell RNA sequencing (scRNA-seq)

Mouse fetal liver and adult mouse bone marrow HSPCs were prepared as a single cell suspension and prepared for FACS as described above. Single cells were index-sorted into 96-well plates containing lysis buffer as described by Liu et al. Briefly, sorted cells were centrifuged at 4C and snap-frozen on dry ice, and stored at −80C immediately after sorting. Reverse transcription (RT) and cDNA pre-amplification were performed using Smart-seq3 protocol with minor modifications. From the purified cDNA, 1uL was taken for quality control for each well. cDNA concentration and size distribution for each well was determined by Fragment Analyzer (Advanced Analytical). Wells with a concentration less than 1.7ng/uL were excluded his cutoff was determined by measuring the concentration in blank wells with ERCC but no sorted cell. Wells with cDNA concentration above the cutoff value were consolidated and reformatted to a new 384-well plate using the Mosquito X1 liquid handler (SPT Labtech), normalizing each well’s concentration to a range of 1.7–4.0 ng/μL by dilution with UltraPure water. Normalized cDNA was used to prepare Illumina sequencing libraries. Tagmentation was performed by combining 0.4 μL cDNA with 1.2 μL homebrew Tn5 mix (1 ng/μL Tn5 enzyme, 16 mM Tris-HCl pH 7.6, 16 mM MgCl2, and 8% dimethylformamide in UltraPure water). The reaction was stopped by adding 0.4 μL neutralization buffer (0.1% SDS). Indexing PCR reactions were performed by adding 0.4 μL of 5 μM i5 indexing primer, 0.4 μL of 5 μM i7 indexing primer (Integrated DNA Technologies, custom made 7680-plex unique dual index-primer set), and 1.2 μL KAPA HiFi HotStart ReadyMix. PCR amplification was performed using the following program: 72°C for 3 min, 95°C for 30 sec, 98°C for 10 sec, 67°C for 30 sec, 72°C for 60 sec, repeating from step 3 for 10 cycles. For each 384-well plate, 1 μL from each well was pooled, followed by purification using 0.8X volume of AMPure XP beads. The 384-cell library pool from each plate was analyzed for concentration and size distribution using a Fragment Analyzer. Twenty 384-cell library pools were normalized for concentration, pooled to form a 7680-plex library pool, purified, and concentrated using 0.8X AMPure beads. The final 7680- plex library pool was sequenced on a NovaSeq 6000 S4 flow cell (Illumina) to obtain ∼1–2 million 2×150 base-pair paired-end reads per cell.

#### Read mapping

Sequences were demultiplexed using bcl2fastq version 2.19.0.316. 3’ adapter sequences were removed from reads using skewer v0.2.2^39^, and aligned to the hg38 genome (Gencode version GRCm38.p6) with STAR aligner version 2.6.1d using 2-pass mapping^40^. Briefly, as a first pass, reads for every cell were aligned using STAR genome index generated using the Gencode transcript annotation for the human genome (version 34). Mapped splice junctions for each cell from the first-pass mapping were extracted, aggregated together and a new STAR index was created where any newly discovered splice junctions were included in addition to the existing Gencode annotation during genome index generation. The new STAR index with all known and newly identified splice-junctions was then used for second pass read mapping. Parameters used for STAR mapping were adapted from the ENCODE long-mRNA-pipeline (https://github.com/ENCODE-DCC/long-read-rna-pipeline) recommendations, also detailed in the STAR manual. In addition to the ENCODE recommended options we also used the “--quantMode TranscriptomeSAM” option during second-pass mapping to generate a bam file containing a catalog of all reads mapped to the transcriptome. This bam file was used as input to calculate expression levels of either genes (sum total of expression levels of all known transcript variants) or individual transcripts using RSEM version 1.3.3 with settings “--single-cell-prior”, which accounts for the sparse nature of mRNA detection usually prevalent in scRNA-seq.

#### Data preprocessing

Gene count tables were combined with metadata using the Scanpy python package v.1.8.2. We filtered out genes expressed in fewer than 3 cells, as well as cells with fewer than 500 detected genes or 5000 read counts. The data were normalized using size factor normalization so that every cell has 10,000 read counts, log transformed and scaled to a maximum value of 10. Highly variable genes were computed using default parameters. We then performed principal component analysis, computed the neighborhood graph, and clustered the data using the Leiden method. PAGA was used to visualize data, as well as reconstruct gene expression changes along maturation trajectories.

### Lentiviral production and transduction of FL HSCs

Lentivirus was produced in HEK293T cells (ATCC, Cat. CRL-3216) maintained in antibiotic-free medium per ATCC guidelines and used below passage 10. Viral particles were generated using the TransIT® Lentiviral Transfection System (Mirus, Cat. MIR 6655) according to the manufacturer’s protocol. Briefly, 10 cm plates were transfected with lentiviral transfer, packaging, and envelope plasmids. Plasmids were propagated using Stable competent cells (NEB, Cat. C3040H) and purified with an EndoFree Plasmid Maxi Kit (Qiagen, Cat. 12362).

Viral supernatants were collected at 48 and 72 hours post-transfection, filtered through 0.45 µm filters, and concentrated using a homemade sterile 4× PEG solution (50% PEG 8000 in Hanks’ Balanced Salt Solution). The virus/PEG mixture was incubated overnight at 4 °C, followed by centrifugation at 3,000 × g for 1 hour at 4 °C. Supernatant was discarded and pellets were resuspended in Opti-MEM (Gibco), snap-frozen, and stored at –80 °C. Viral titer was measured using the Lenti-X™ p24 Rapid Titer Kit (Takara, Cat. 631476).

Sorted *Hoxb5* NEO1 or NEO1 HSCs from E14.5 fetal livers were transduced separately with lentiviral vectors encoding membrane-localized fluorescent reporters mKOk or mUKG, respectively. Transductions were carried out in HemeEx media supplemented with 5 µg/mL Polybrene (VectorBuilder, Cat. PL200), 10 ng/mL SCF (Peprotech, Cat. AF-250-03), and 100 ng/mL TPO (Peprotech, Cat. AF-315-14). Cells were incubated for 24 hours in low-oxygen conditions (5% CO, 5% O) prior to transplantation.

### In utero FL HSC transplantation

CD45.2^+^ E12.5 Hoxb5 fetal liver HSCs (Lineage Sca1 c-Kit CD48 CD150 mCherry^+^) were sorted and further sorted by NEO1 or NEO1 expression. NEO1 HSCs were lentivirally transduced with membrane-localized mKOk, and NEO1 HSCs with mUKG, as described above.

Equal numbers of NEO1 (mKOk) and NEO1 (mUkG) HSCs were mixed and co-transplanted into the fetal liver of E12.5 CD45.1 PepBoy embryos. Pregnant dams were anesthetized with isoflurane, and a midline laparotomy was performed to expose the uterine horns. Each embryo received a 5 µL injection of the HSC mixture in sterile saline, delivered directly into the fetal liver using a 34G sub-microliter syringe (NANOFIL, World Precision Instruments). After transplantation, the uterus was returned to the abdominal cavity and the dam was sutured and allowed to recover.

Embryos were harvested at E18.5, and fetal livers were collected for flow cytometric analysis. Donor-derived cells were identified as CD45.2 CD45.1 and further distinguished by expression of mKOk or mUkG. The contribution of each donor population to the HSC compartment (Lineage Sca1 c-Kit CD48 CD150) and differentiated progeny was quantified to assess relative contribution and lineage potential.

### Quantification and statistical analysis

For all experiments, *n* indicates the number of biologically independent replicates and are indicated in the figure legends. All statistical analysis was performed using GraphPad Prism (GraphPad Software) unless otherwise noted. The statistical tests used (parametric or nonparametric *t*-tests) are noted in the figure legends. Error bars denote mean ± S.E.M.

## Acknowledgements

We thank the members of the Weissman lab for advice and discussions; Linda Quinn and Teja Naik for laboratory management; Tal Raveh for assistance with general grant administration and funding acquisition. Aaron McCarty and Charlene Wang for mouse colony management; Catherine Carswell-Crumpton, Cheng Pan, and Joe Pasillas for FACS support. This work was supported by the NIA R36 award R36-AG090859 to A.B, the Stanford University Dean’s Fellowship and the Walter V. and Idun Berry Postdoctoral Fellowship to E.K-S, the ETIUDA grant from the Polish National Science Centre no. UMO-2019/32/T/NZ3/00624 to M.Z, the TL1DK139565 to A.T.B, the NIH/NCI Outstanding Investigator Award R35-CA220434 to I.L.W, and the Virginia and D.K. Ludwig Fund for Cancer Research to I.L.W.

## Author contributions

A.B. conceived the study, designed and performed all experiments, analyzed data, and wrote the manuscript. M.B. and L.Y. performed experiments and analyzed data with A.B. E.K- S. and M.Z. designed and performed experiments. A.T.B., L.S.W., U.L., A.Z., N.G., B.Z., N.W.G., R.H. and R.S. assisted with experiments and sample preparation. B.O.G. performed sample processing for SMART-seq3 single-cell RNA sequencing. I.L.W. conceived, supervised the study, and edited the manuscript. A.B., M.B., and L.Y. contributed equally to this work as co-first authors.

