## Supplemental for "Neogenin-1 marks myeloid-primed fetal hematopoietic stem cells that undergo progressive lineage-restriction with age"

Supplementary Figure 1

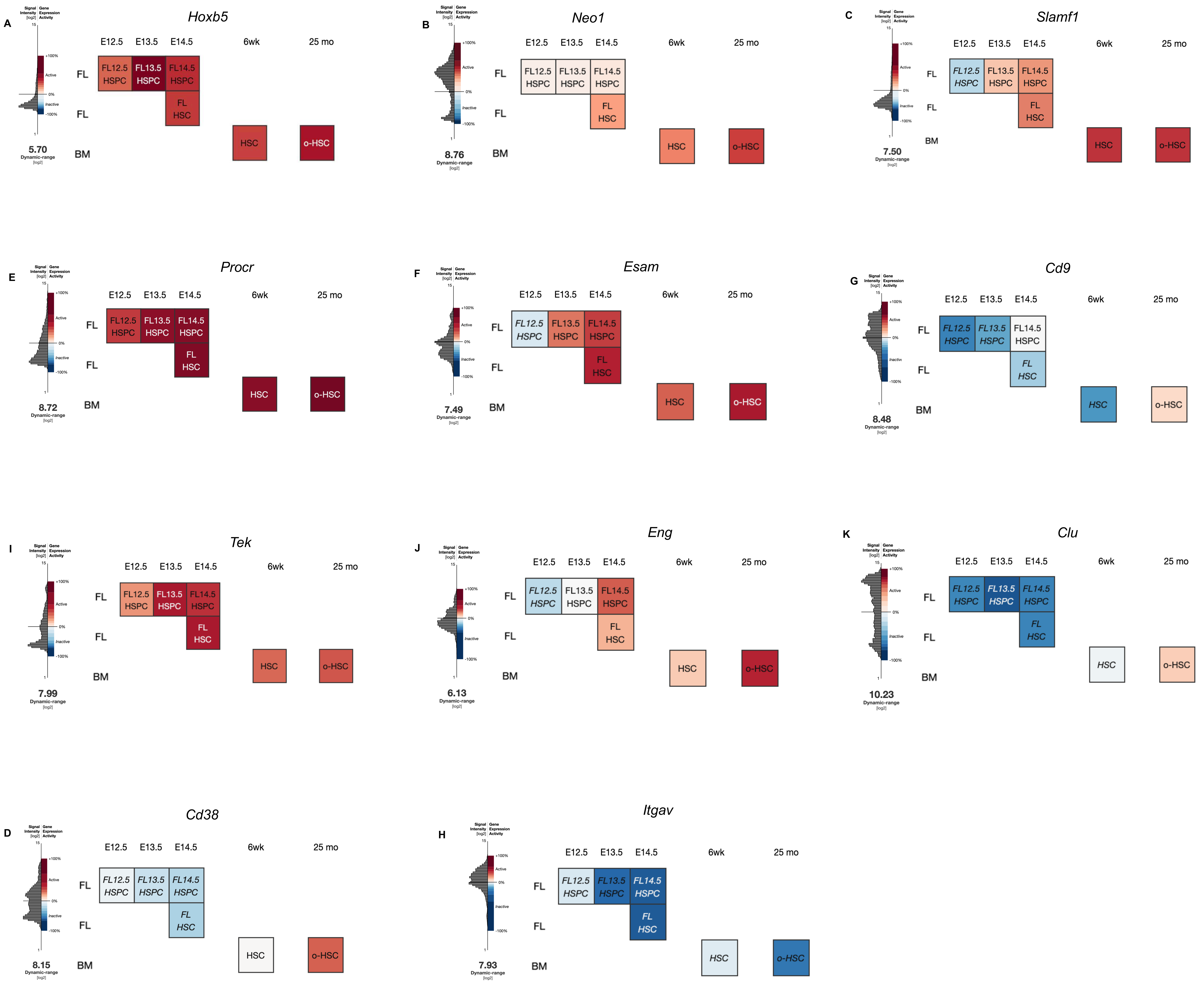

Supplementary Figure 2

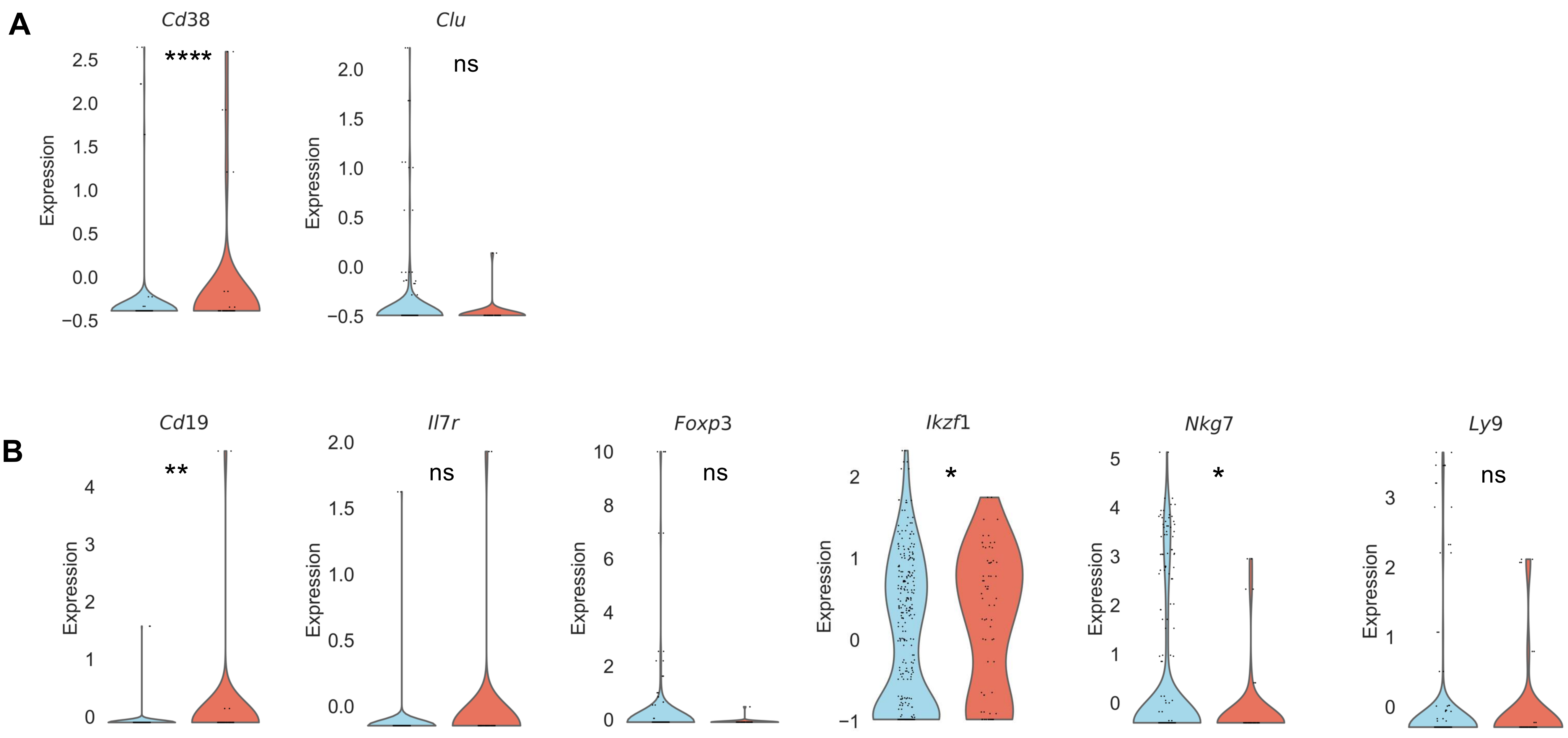

Supplementary Figure 3

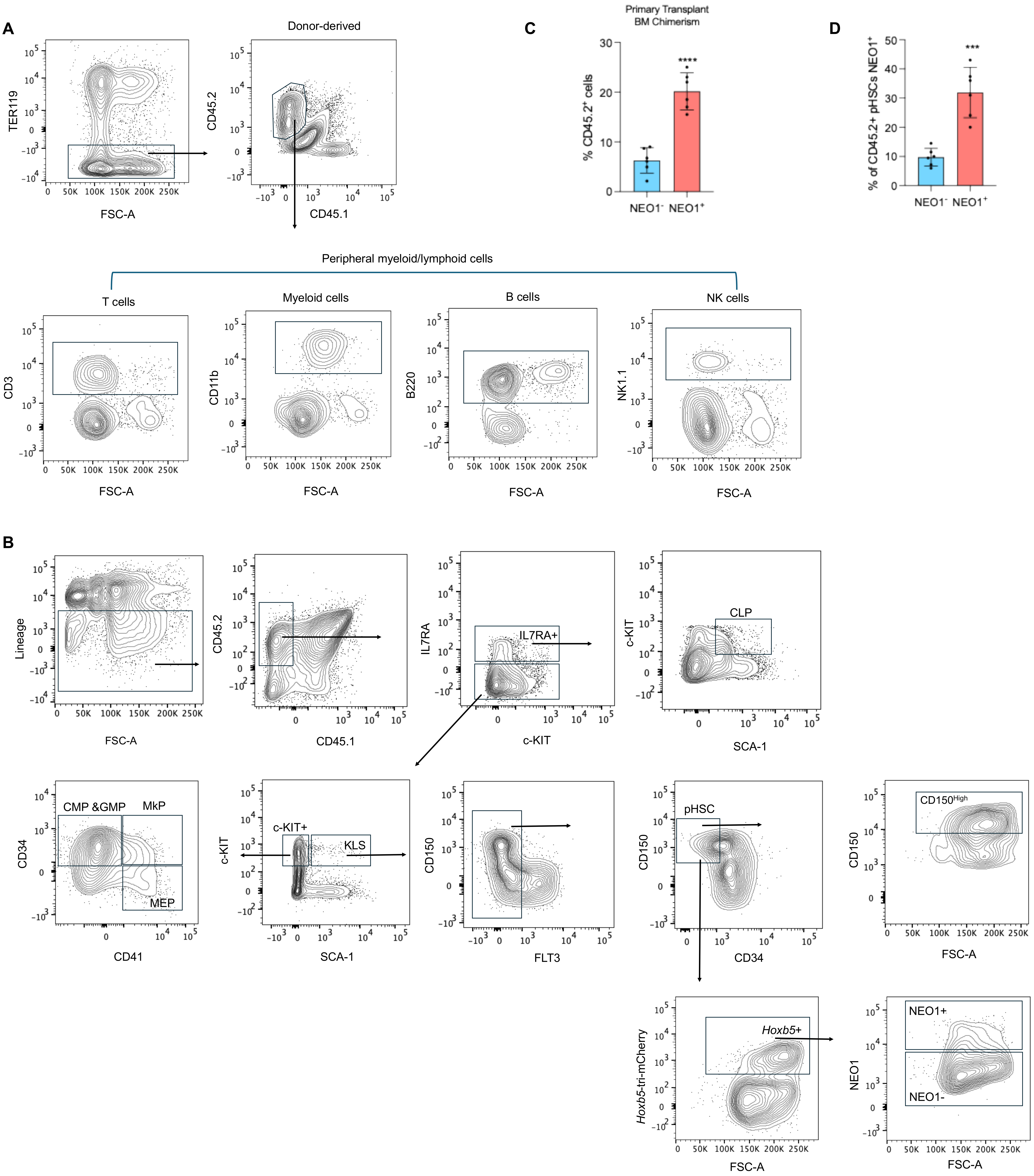

Supplementary Figure 4

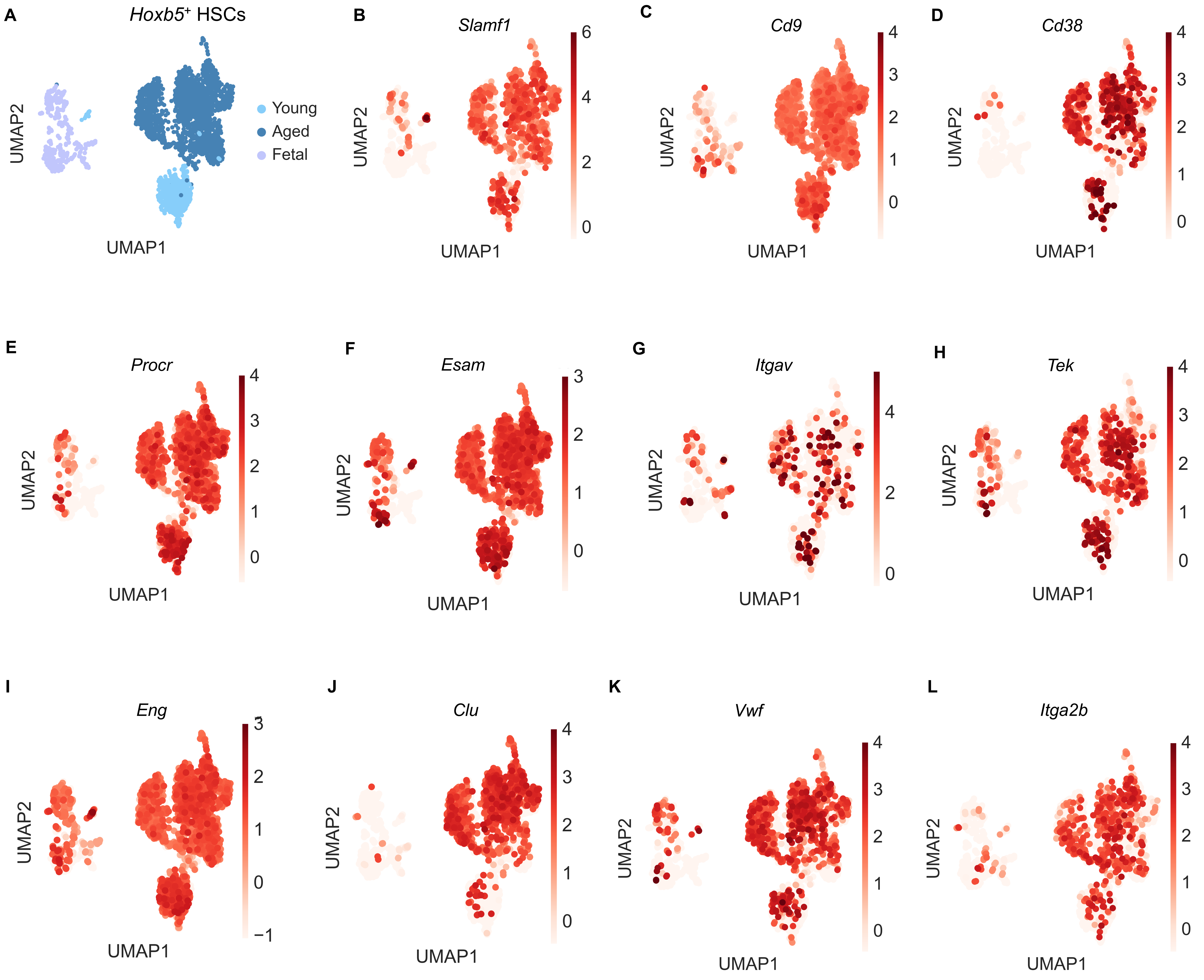

Supplementary Figure 5

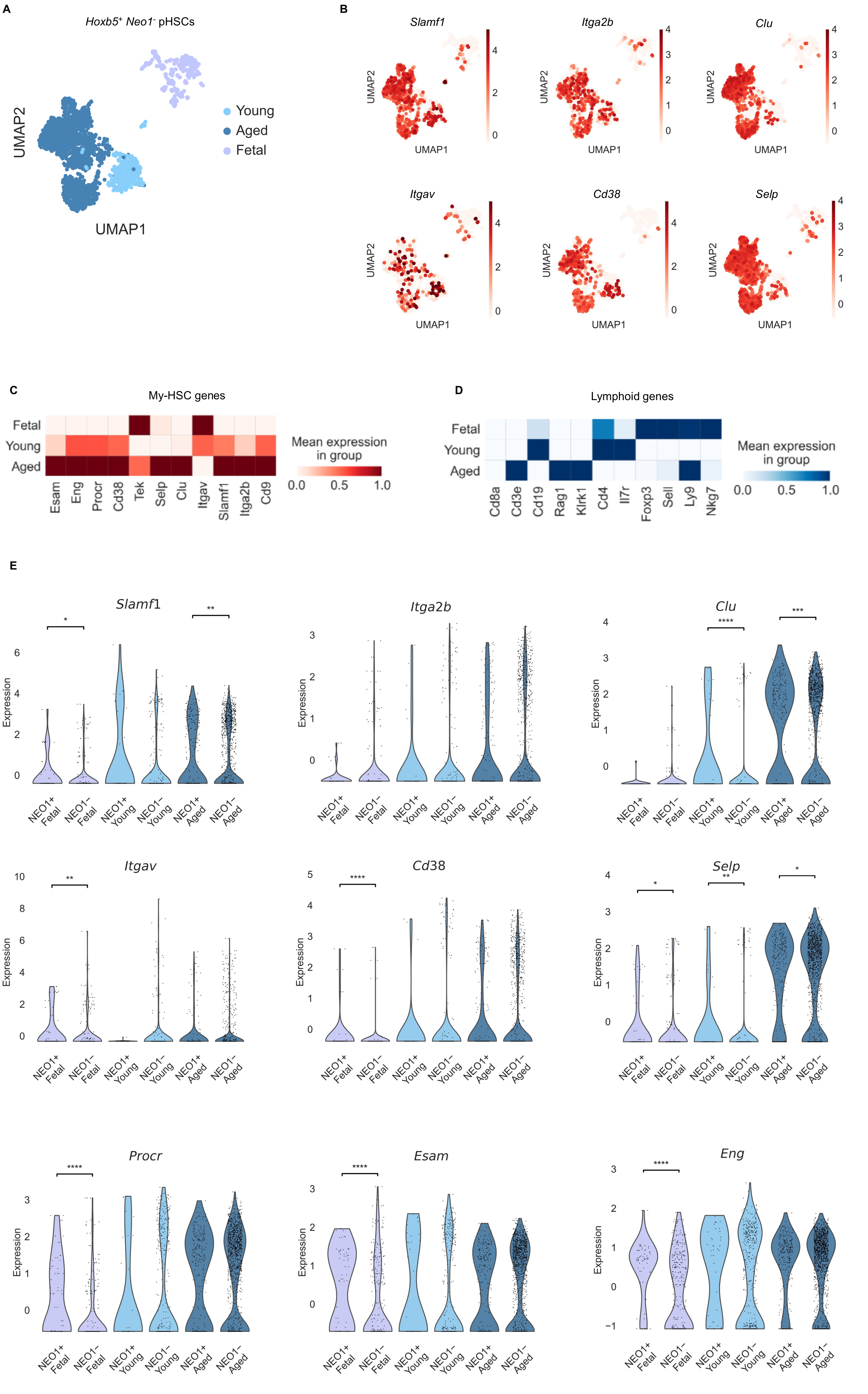

Supplementary Figure 6

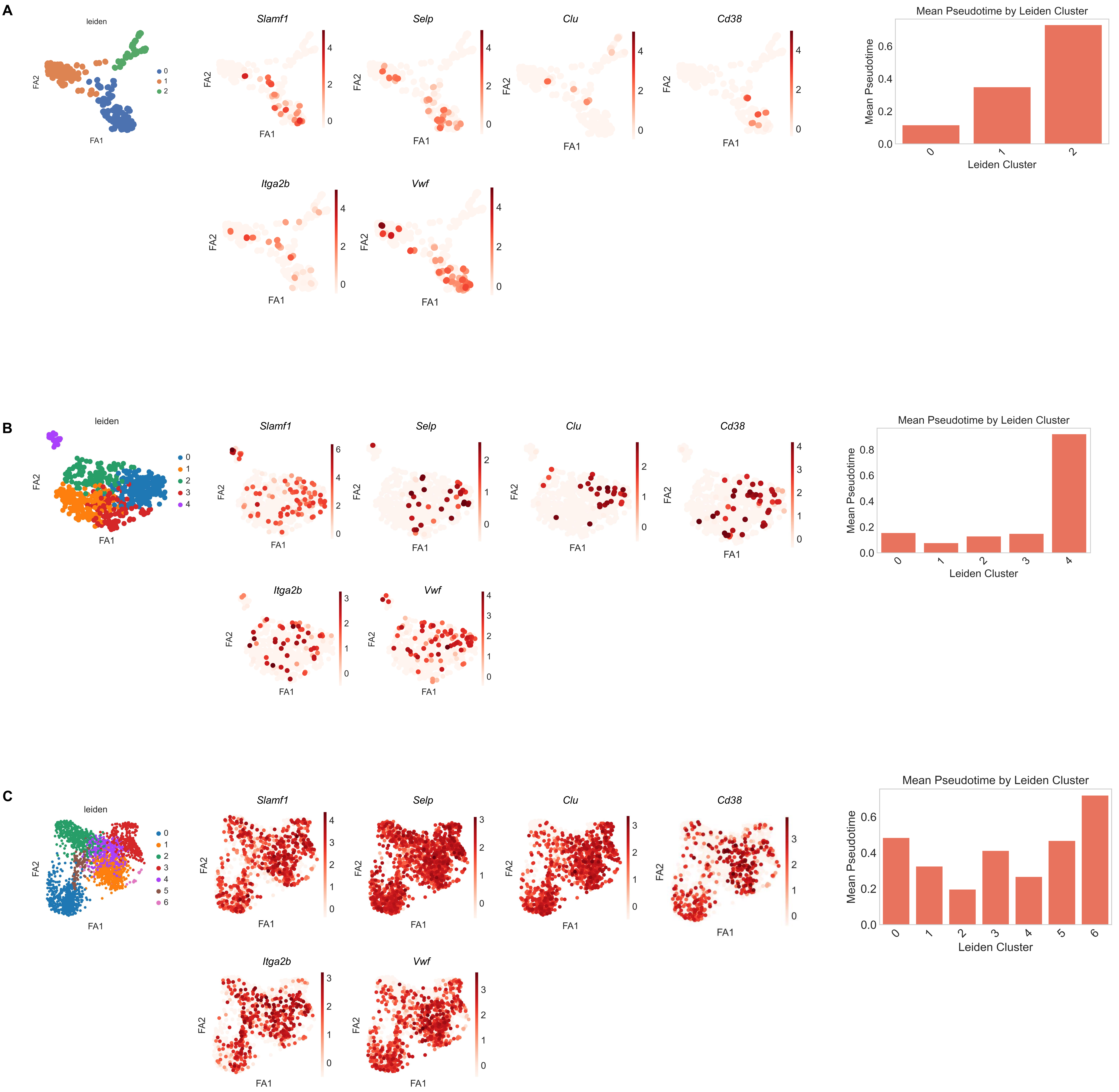
